# Versatile mapping-by-sequencing with Easymap v.2

**DOI:** 10.1101/2022.07.14.500089

**Authors:** Samuel Daniel Lup, Carla Navarro-Quiles, José Luis Micol

**Affiliations:** Instituto de Bioingeniería, Universidad Miguel Hernández, Campus de Elche, 03202 Elche, Spain

**Keywords:** mapping-by-sequencing, candidate mutations, forward genetics, NGS, variant density mapping, serial backcrossing, QTL-seq

## Abstract

**Motivation:** Mapping-by-sequencing combines Next Generation Sequencing (NGS) with classical genetic mapping by linkage analysis to establish gene-to-phenotype relationships. Although numerous tools have been developed to analyze NGS datasets, only a few are available for mapping-by-sequencing. One such tool is Easymap, a versatile, easy-to-use package that performs automated mapping of point mutations and small insertion/deletions (InDels), as well as large DNA insertions.

**Results:** Here, we describe Easymap v.2, which includes additional workflows to perform QTL-seq and variant density mapping analyses. Each mapping workflow can accommodate different experimental designs, including outcrossing and backcrossing, F_2_, M_2_, and M_3_ mapping populations, chemically induced mutation and natural variant mapping, input files containing single-end or paired-end reads of genomic or complementary DNA sequences, and alternative control sample files in FASTQ and VCF formats. Easymap v.2 can also be used as a variant analyzer in the absence of a mapping algorithm and includes a multi-threading option.

**Availability and implementation:** Code is available at http://genetics.edu.umh.es/resources/easymap/

## INTRODUCTION

Identifying the causal genetic variant for a phenotype of interest is a common starting point in the genetic dissection of a biological process. Individuals exhibiting a phenotype of interest can be isolated by screening a large set of wild-type accessions or natural races or by the mutagenesis of a wild-type strain to isolate phenotypically distinct mutants among its progeny. The commonly used mutagen ethyl methanesulfonate (EMS) induces point mutations (usually G→A transitions) in random positions across the genome, some of which alter the sequence of genes and/or their transcriptional or post-transcriptional regulation (James and Dooner, 1990; Jansen *et al.*, 1997).

A classic approach to mapping the causal mutation is linkage analysis between the mutation and molecular markers in segregating populations. This procedure has been integrated with Next Generation Sequencing (NGS): the improved technique is known as “mapping-by-sequencing” (Candela *et al.*, 2015; Hartwig *et al.*, 2012; James *et al.*, 2013; Schneeberger and Weigel, 2011). In a typical mapping-by-sequencing experiment, the distribution of allele frequencies of biallelic Single Nucleotide Polymorphisms (SNPs) is studied in a mapping population: a pool of phenotypically recessive mutant individuals selected from a segregating population. The mapping population is used to identify genomic regions where SNP allele frequency is influenced by the phenotypic selection performed (James *et al.*, 2013; Schneeberger *et al.*, 2009; Wachsman *et al.*, 2017). In the model plant Arabidopsis (*Arabidopsis thaliana*), bulked segregant analysis is usually (but not exclusively) performed using populations composed of F_2_ individuals generated from the selfing of an F_1_ progeny derived from a cross between a mutant and a wild-type strain. The mutant can be crossed to a genetically divergent from—and hence polymorphic to—its pre-mutagenesis wild-type parent (outcross or map cross), or to the wild-type parent itself (backcross or isogenic cross).

Another common approach to uncovering gene-to-phenotype relationships is to identify genetic lesions in a population of phenotypically mutant individuals obtained from recurrent backcrosses to a reference strain (Doitsidou *et al.*, 2016; Klein *et al.*, 2018). This approach, which was first used to identify EMS-induced mutations, is called EMS variant density mapping (Minevich *et al.*, 2012; Zuryn *et al.*, 2010). This technique relies on the presence or absence of variants along the genome and the detection of genomic regions with a significantly higher density of variants (high-density variant peaks or clusters) compared to the rest of the genome. These regions, which show linkage disequilibrium, are expected to contain the mutation causing the phenotype of interest, along with a set of tightly linked variants selected through recurrent backcrossing. This mapping strategy is convenient when selecting numerous mutants from a segregating population is not feasible due to complex or expensive phenotyping, scarce offspring, or life cycles that hinder the isolation of recombinant individuals. This approach is however slower than conventional mapping-by-sequencing strategies, since several backcrosses are needed to obtain the mapping population (Table 1). There are currently no user-friendly, graphic interface-based bioinformatic tools that automate the analysis of datasets obtained from serial backcrossing mapping strategies.

**Table 1.**
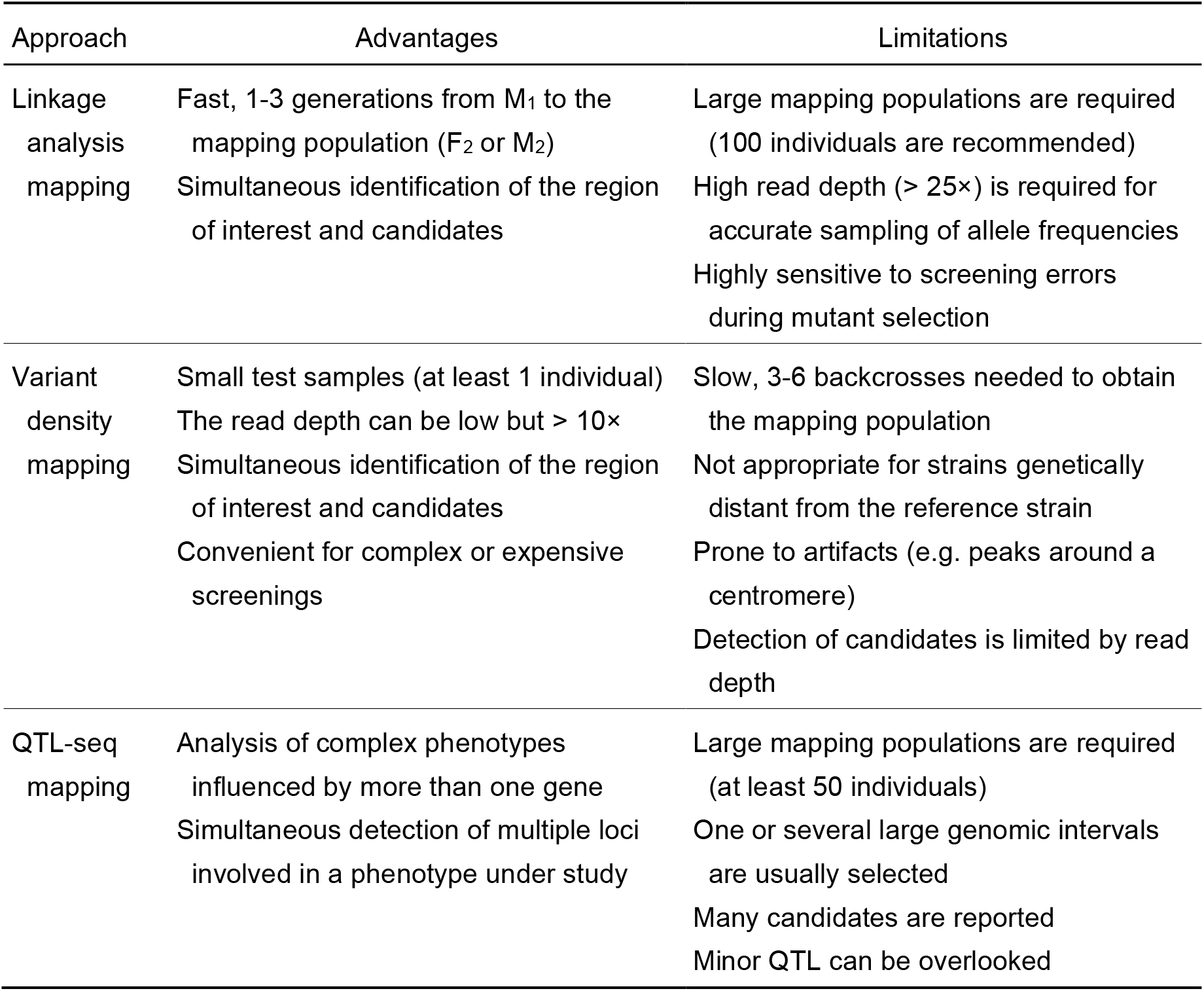
Experimental approaches for mapping-by-sequencing of SNPs and small InDels

| Approach | Advantages | Limitations |
| --- | --- | --- |
| Linkage analysis mapping | Fast, 1-3 generations from M<sub>1</sub> to the mapping population (F<sub>2</sub> or M<sub>2</sub>)<br>Simultaneous identification of the region of interest and candidates | Large mapping populations are required (100 individuals are recommended)<br>High read depth (> 25×) is required for accurate sampling of allele frequencies<br>Highly sensitive to screening errors during mutant selection |
| Variant density mapping | Small test samples (at least 1 individual)<br>The read depth can be low but > 10×<br>Simultaneous identification of the region of interest and candidates<br>Convenient for complex or expensive screenings | Slow, 3-6 backcrosses needed to obtain the mapping population<br>Not appropriate for strains genetically distant from the reference strain<br>Prone to artifacts (e.g. peaks around a centromere)<br>Detection of candidates is limited by read depth |
| QTL-seq mapping | Analysis of complex phenotypes influenced by more than one gene<br>Simultaneous detection of multiple loci involved in a phenotype under study | Large mapping populations are required (at least 50 individuals)<br>One or several large genomic intervals are usually selected<br>Many candidates are reported<br>Minor QTL can be overlooked |

Most phenotypic traits are influenced by multiple genes and their interactions with the environment. Quantitative trait loci (QTL) are genomic regions containing genes that contribute to a specific quantitative phenotype, which in plants include agronomically relevant traits such as plant height, biomass production, and pathogen resistance (Alonso-Blanco and Koornneef, 2000; Kearsey, 1998; Kearsey and Farquhar, 1998). QTL were traditionally mapped by linkage analysis in the segregating progeny of a cross of two strains that genetically differ for a quantitative trait of interest (Chen *et al.*, 2021; Juenger *et al.*, 2005). This approach was combined with NGS to create QTL-seq, a technique involving the sequencing of two pools of individuals with opposite phenotypes selected from a population that segregates for a number of genetic variants (Takagi *et al.*, 2013). QTL-seq can be used to identify linkage disequilibrium in genomic regions that potentially contain QTL for the trait under study.However, only a few tools have been developed for the analysis of QTL-seq datasets, and these tools require the use of additional software, thus creating complex bioinformatic pipelines (Mansfeld and Grumet, 2018; Wu *et al.*, 2019).

Easymap was developed as a user-friendly software package to facilitate conventional mapping-by-sequencing of point mutations and tagged-sequence mapping of large insertions, both using NGS datasets (Lup *et al.*, 2021). Easymap implements mapping workflows for diverse types of datasets, including DNA whole-genome resequencing and transcriptome sequencing (RNA-seq) data, mapping populations obtained by backcrossing, outcrossing or selfing of a mutant, and control samples consisting of the whole-genome sequences of any parental line of the mapping population or a pool of phenotypically wild-type siblings of the mapping population. Here, we describe Easymap v.2, an updated version of Easymap that features variant density and QTL-seq mapping workflows to detect any spontaneous or mutagen-induced SNPs and small insertion/deletions (InDels), which we refer to collectively here as variants. Easymap v.2 also includes a variant analyzer to explore the effects of a list of variants on genes that contain these variants and on their products. In addition, Easymap v.2 contains a preprocessing module for FASTQ files, supports the use of Variant Call Format (VCF) files as control samples, and allows multithreading. Easymap v.2 is open source and available for download at http://genetics.edu.umh.es/resources/easymap/. We recommend the Quickstart Installation Guide, which any person with no bioinformatics skills can follow to install a fully functional Easymap v.2 program.

## METHODS

### Architecture

Easymap v.2 works in the Unix-based operating systems Ubuntu, Red Hat, Fedora and AMI. It can also be used in Windows 10 within the Ubuntu apps currently available at Microsoft and in virtual machines running a Unix-based operating system within macOS. Easymap v.2 can also be installed and accessed remotely (e.g., in a computational cluster or the Amazon Elastic Compute Cloud service) through its graphical and command line interfaces.

The installation of Easymap v.2 is automated, with a single script that compiles and installs all required software and third-party tools: Python2 (https://www.python.org/about/), Python Imaging Library (https://pillow.readthedocs.io/en/stable/), Virtualenv (https://virtualenv.pypa.io/en/latest/), HTSlib (http://www.htslib.org/), HISAT2 (Kim *et al.*, 2019), Bowtie2 (Langmead and Salzberg, 2012), SAMtools (Li *et al.*, 2009), and BCFtools (Narasimhan *et al.*, 2016).

The installation script also launches the graphical web interface once installation is complete. The Easymap v.2 Quickstart Installation Guide (Supplementary File 1) provides detailed information about how to install Easymap v.2 without any prior bioinformatics knowledge. Advanced installation setups and usage instructions can be found in the Easymap v.2 Documentation (Supplementary File 2).

### Testing

Easymap v.2 was tested on regular desktop computers and on high-performance machines, performance depends on the machine being used and the computational resources allocated to the program. For example, a typical linkage analysis from an Arabidopsis (genome size of ~135 Mb; The Arabidopsis Genome Initiative, 2000) mapping population derived from a backcross, in which test and control samples have a read depth of 50×, can take 6-8 hours using a standard computer without multi-threading. However, the same analysis involving larger genomes such those of maize (*Zea mays*, ~2.4 Gb; Haberer *et al.*, 2005) and barley (*Hordeum vulgare*, ~5.3 Gb; The International Barley Genome Sequencing Consortium, 2012) can take weeks. Therefore, multi-threading is highly recommended when working with large genomes or with experimental designs involving an outcross and can easily be set up using the graphic interface. Easymap v.2 also allows multiple projects to be executed simultaneously, but this can reduce the overall performance of a desktop computer. A minimum of 8 Gb of RAM and available disk storage at least twice the size of all input reads (or three-times the size if pre-processing is enabled) should suffice for most analyses.

## RESULTS

### Variant density mapping workflow

We implemented a workflow in Easymap v.2 that performs variant density mapping in a test sample (Figure 1). The test sample consists of NGS reads obtained from a pool of individuals exhibiting a phenotype of interest that were subjected to several (usually 3 to 6) backcrosses to the reference strain. The use of a control sample is strongly advised. The control sample consists in reads obtained from an individual (or pool of individuals) that shares a considerable number of variants with the test sample. These variants are not related to the phenotype of interest and therefore must be filtered out from the test sample to aid in the identification of high-density variant peaks and candidate variants. In this manner, control reads can be obtained from strains that do not show the phenotype of interest but are genetically related to the test strain, such as the pre-mutagenesis wild-type strain, the parental reference strain, phenotypically wild-type siblings of the mapping population, or other mutant lines isolated from the same mutagenesis screen (Figure 1A).

**Figure 1.**
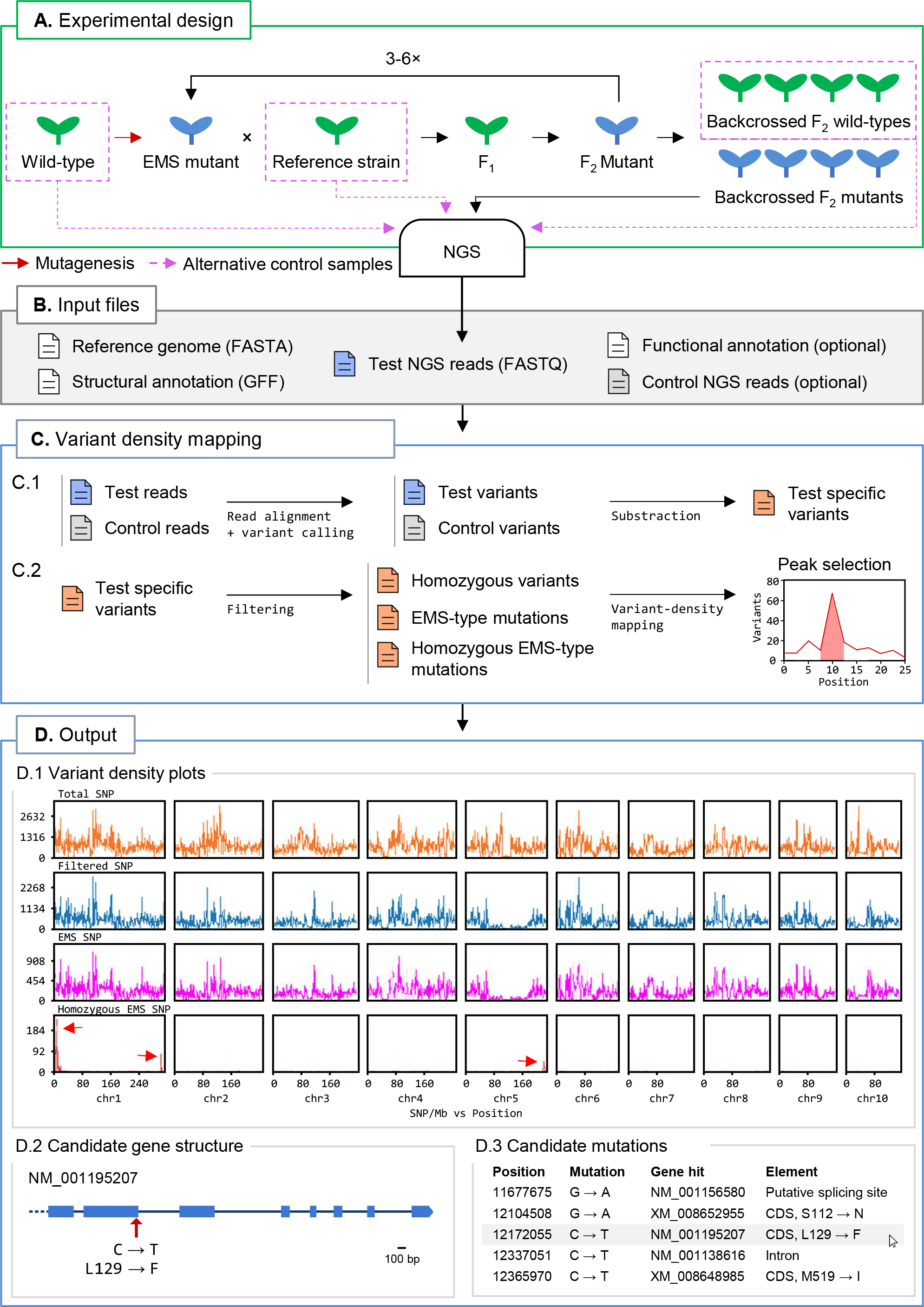
Variant density mapping with Easymap v.2. (A) Overview of the experimental design. A mutant of interest (in blue) carrying an EMS-induced mutation that is causal for a target trait is backcrossed 3 to 6 times to its reference strain to generate a test sample of backcrossed F_2_ mutants. DNA extracted from a pool of mutants and a control sample is subjected to NGS to obtain the test and control reads. (B) Input files. A FASTA file with the reference genome sequence, the corresponding GFF file with structural annotation of the genome, and the test NGS reads are required. A read file from a control sample containing genetic variants not linked to the causal mutations is strongly recommended. A functional annotation file is optional. (C) Easymap v.2 variant density mapping workflow. Files are color-coded in blue, green and red for the test, control and test-specific (filtered) variants, respectively. The arrows represent steps of the analysis performed with third-party software (alignment and variant-calling) and proprietary Python scripts. (C.1) The test and control reads are aligned to the reference genome to detect variants that distinguish each sample from the reference sequence. The control variants are then subtracted from the test variants to obtain the test-specific variants. (C.2) A series of filtering steps generates the lists of homozygous variants, EMS-type mutations, and homozygous EMS-type mutations, which are used to detect high-density peaks of variants, generate plots, and extract candidate variants. (D) Easymap v.2 output obtained from variant density mapping analysis. (D.1) Plots of the number of total, test-specific, EMS-type, and homozygous EMS-type variants per 1-Mb bin. The red arrows point to peaks of EMS-type variants in the maize genome, the first of which contains the causal mutation *teosinte branched1 enhancer* (*ten*) (Klein *et al.*, 2018). The two other arrows point to random artifacts found in the original publication as well. (D.2) Structure of the NM_001195207 transcription unit. The red arrow indicates the position of the ten mutation. Equivalent diagrams are generated for each candidate gene. (D.3) An extract of the list of candidate genes, with the *ten* gene highlighted.

Once the input files (comprising the test and control reads) have been loaded by the user (Figure 1B), Easymap v.2 reports the list of test sample-specific variants. This list is used to generate two sublists: one containing homozygous variants, and the other all EMS-type mutations. A third sublist that contains the homozygous EMS-type variants is created by the intersection of the first two sublists (Figure 1C). Easymap v.2 then detects high-density variant peaks along the genome of the test sample in overlapping sliding windows and establishes regions of interest according to the variant density distribution (Figure 1D.1). The variants within the regions of interest are reported as candidate mutations if they are located within a gene (Figure 1D.2 and D.3). In the web interface, Easymap v.2 provides diagrams representing each gene of interest, plots of the distribution of variant density along the genome, and a table listing extensive information about each variant.

To test the functionality of the variant density mapping workflow, we reproduced results from nine previously published datasets, including studies in the nematode *Caenorhabditis elegans* and maize, and detected the known causal mutation in all instances (Table 2). The datasets from mutants in the reference background (Svensk *et al.*, 2016; Zuryn *et al.*, 2010) provided fairly clear information, as the number of background variants was limited, resulting in a generally approachable number of candidate causal mutations.

**Table 2.** Validation of the Easymap v.2 variant density mapping workflow with experimental data

| Original study | Species | Mutant | Variants identified by EasyMap v.2 |
| --- | --- | --- | --- |
| Zuryn *et al.* (2010) | *C. elegans* | *mutA* | The causal mutation and two other nonsynonymous mutations in the candidate region |
|  |  | *mutD* | The correct candidate region (the causal mutation was unknown) |
|  |  | *mutH* | The correct candidate region (the causal mutation was unknown) |
| Zuryn *et al.* (2014) | *C. elegans* | *jmjd-3.1* | The causal mutation was the only nonsynonymous mutation in the candidate region |
|  |  | *egl-27* | The causal mutation and two other nonsynonymous mutations in the candidate region |
| Svensk *et al.* (2016) | *C. elegans* | *sma-1* | The causal mutation and eight other nonsynonymous mutations in the candidate region |
|  |  | *dpy-23* | The causal mutation and ten other nonsynonymous mutations in the candidate region |
|  |  | *sma-9* | The causal mutation and another nonsynonymous mutation in three candidate regions |
| Klein *et al.* (2018) | *Z. mays* | *ten* | The causal mutation and 249 other nonsynonymous mutations in the candidate region |

Further information is shown in Supplementary Table S1.

The use of datasets generated to map mutations in a background that is genetically distant from that of the reference strain (Klein *et al.*, 2018) generally results in larger numbers of candidates due to the high density of natural polymorphisms between the two strains. In general, additional fine-mapping experiments are needed to identify the causal mutation.

### QTL-seq mapping workflow

Another workflow implemented in Easymap v.2 performs QTL-seq mapping analysis from two pools of individuals of a given segregating population with opposite phenotypes (Figure 2A). After loading the input files (Figure 2B), the QTL-seq mapping workflow uses SNPs common to both pools to identify the differences between the allele frequencies of each sample (dAF) in sliding windows across the genome (Figure 2C). This step allows the software to select genomic regions in which the dAF deviates from 0, i.e., there is opposite linkage disequilibrium in both samples. The selected regions are reported as potential QTL that contain candidate variants and genes, and a set of figures and tabular data is generated to allow the user to consider whether these candidates are modifiers of the phenotype under study (Figure 2D). As QTL-seq is a common approach for characterizing agronomically relevant traits in cultivars and species that lack a proper structural annotation of the genome, we enabled the possibility to run the QTL-seq mapping workflow without a structural annotation file (usually in genome feature file [GFF] format). Without a GFF file, this workflow can identify candidate regions that might contain QTL, but gene annotations and identification will not be available in the report.

**Figure 2.**
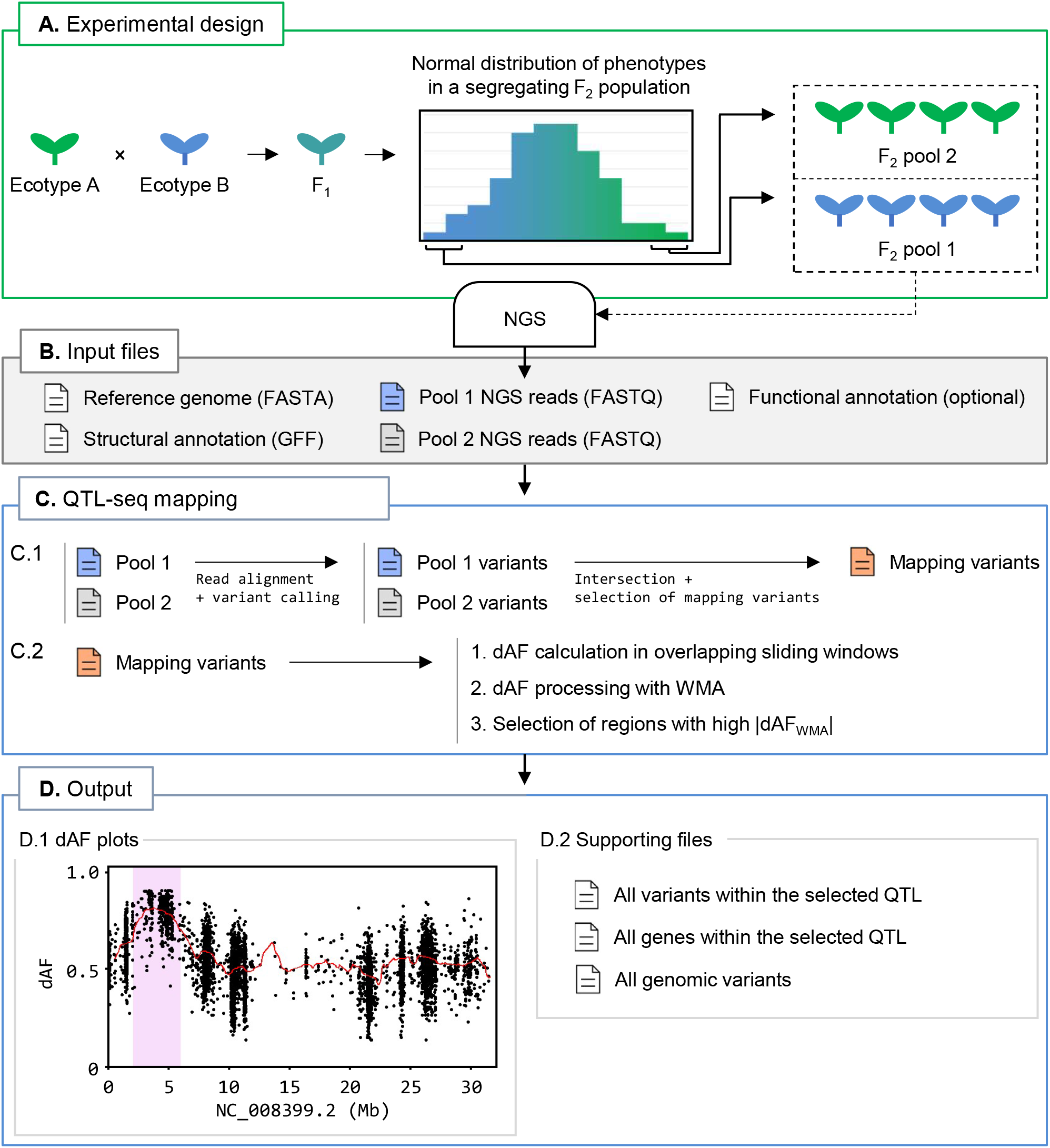
QTL-seq mapping with Easymap v.2. (A) Overview of the experimental design. Two wild-type accessions that genetically differ for a quantitative trait of interest (in blue and green) are crossed. The F_1_ progeny is selfed to generate a segregating F_2_ population. Plants exhibiting the most extreme phenotypes for the trait of interest are bulked into two pools for DNA extraction and NGS. (B) Input files for Easymap v.2. Files containing the reference genome sequence and the NGS reads from both pools mentioned above are required. Structural and functional annotation files are optional. (C) Easymap v.2 QTL-seq mapping workflow. Files are color-coded in blue, green and red for pools 1 and 2, and mapping variants, respectively. The arrows represent steps performed with third party software (alignment and variant-calling) and proprietary Python scripts (all remaining steps). (C.1) Reads from pools 1 and 2 are processed to generate variant files, which are intersected. Segregating variants that are present and have an allele frequency lower than 1 in both samples are then selected to create a list of mapping variants. This step filters out variants common to both parental lines. (C.2) The difference between the allele frequency values between the mapping variants of the two samples (dAF) are calculated and averaged in overlapping sliding windows. dAF values per window are then averaged with the values of adjacent windows via weighted moving averages (WMA) to smooth out the mapping signal, generating dAF_WMA_ values. Finally, the processed dAF_WMA_ values are searched for regions with nonzero values to select those positively or negatively influenced by phenotypic selection. (D) Easymap v.2 output from a QTL-seq mapping experiment in rice (Takagi *et al.*, 2013). (D.1) dAF plotted along a chromosome containing a candidate QTL, highlighted in pink. The mapping variants are represented by black dots. Processed dAF_WMA_ values are represented by a red line. These plots are generated for each chromosome. (D.2) Supporting files are provided to assess the results of mapping analysis and to establish alternative or additional regions of interest. Diagrams of the candidate genes and tables including the candidate variants and genes are reported when a GFF file is provided.

To test the QTL-seq mapping workflow, we reproduced results from 14 different QTL-seq analyses in tomato (*Solanum lycopersicum*), barley, and different rice (*Oryza sativa*) cultivars using F_2_ (Illa-Berenguer *et al.*, 2015; Takagi *et al.*, 2013; Wang *et al.*, 2018; Yang *et al.*, 2017), M3 (Fekih *et al.*, 2013), double haploid (Hisano *et al.*, 2017), and Recombinant Inbred Line (RIL; (Fekih *et al.*, 2013) mapping populations. These datasets included whole-genome and exome sequencing datasets, some with suboptimal average read depths (below 8×; Table 3). Data analysis and criteria for QTL selection varied markedly among these studies. To provide robust results, Easymap v.2 performs a stringent selection of mapping variants for the detection of major QTL. However, we recommend that the user inspect the dAF plots, as well as the supporting files produced by Easymap v.2, to detect additional regions of interest that might have been overlooked, such as minor QTL. In our validation experiments, major QTL were selected correctly, but a few minor QTL were missed by Easymap v.2. These minor QTL became evident after visual inspection of the final report produced by our software. Identification of the variants that affect the phenotype under study is restricted by the availability of a structural annotation file, as well as the read depth of the dataset. Nonetheless, Easymap v.2 was successful in detecting all previously reported variants in the tested datasets (Hisano *et al.*, 2017; Wang *et al.*, 2018).

**Table 3.**
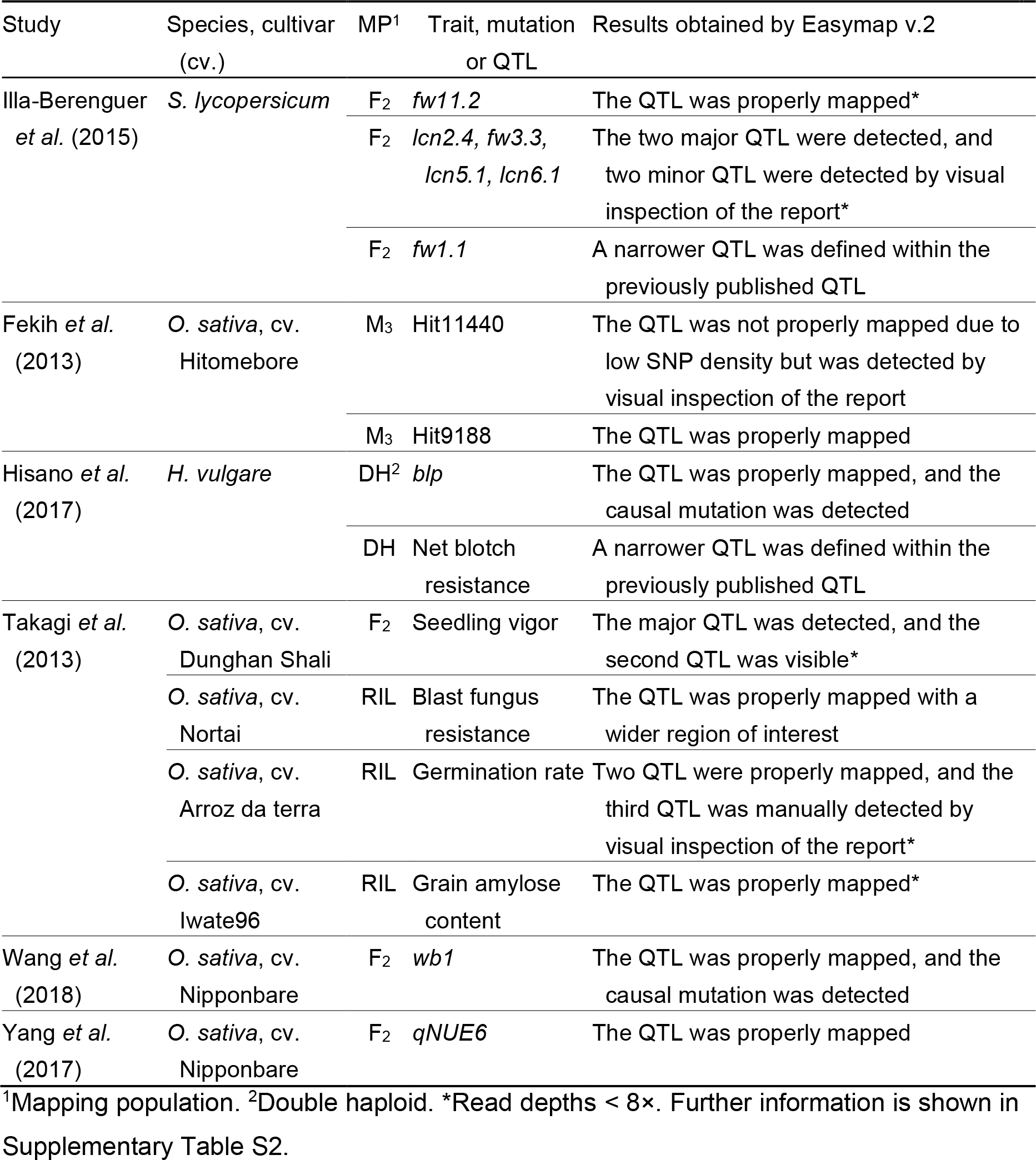
Validation of the Easymap v.2 QTL-seq workflow with experimental data

| Study | Species, cultivar (cv.) | MP<sup>1</sup> | Trait, mutation or QTL | Results obtained by Easymap v.2 |
| --- | --- | --- | --- | --- |
| Illa-Berenguer *et al.* (2015) | *S. lycopersicum* | F<sub>2</sub> | *fw11.2* | The QTL was properly mapped* |
|  |  | F<sub>2</sub> | *lcn2.4, fw3.3, lcn5.1, lcn6.1* | The two major QTL were detected, and two minor QTL were detected by visual inspection of the report* |
|  |  | F<sub>2</sub> | *fw1.1* | A narrower QTL was defined within the previously published QTL |
| Fekih *et al.* (2013) | *O. sativa*, cv. Hitomebore | M<sub>3</sub> | Hit11440 | The QTL was not properly mapped due to low SNP density but was detected by visual inspection of the report |
|  |  | M<sub>3</sub> | Hit9188 | The QTL was properly mapped |
| Hisano *et al.* (2017) | *H. vulgare* | DH<sup>2</sup> | *blp* | The QTL was properly mapped, and the causal mutation was detected |
|  |  | DH | Net blotch resistance | A narrower QTL was defined within the previously published QTL |
| Takagi *et al.* (2013) | *O. sativa*, cv. Dunghan Shali | F<sub>2</sub> | Seedling vigor | The major QTL was detected, and the second QTL was visible* |
|  | *O. sativa*, cv. Nortai | RIL | Blast fungus resistance | The QTL was properly mapped with a wider region of interest |
|  | *O. sativa*, cv. Arroz da terra | RIL | Germination rate | Two QTL were properly mapped, and the third QTL was manually detected by visual inspection of the report* |
|  | *O. sativa*, cv. Iwate96 | RIL | Grain amylose content | The QTL was properly mapped* |
| Wang *et al.* (2018) | *O. sativa*, cv. Nipponbare | F<sub>2</sub> | *wb1* | The QTL was properly mapped, and the causal mutation was detected |
| Yang *et al.* (2017) | *O. sativa*, cv. Nipponbare | F<sub>2</sub> | *qNUE6* | The QTL was properly mapped |

<sup>1</sup>Mapping population. <sup>2</sup>Double haploid. \*Read depths < 8x. Further information is shown in Supplementary Table S2.

### Additional implementations

#### Variant analyzer workflow

Easymap v.2 includes a variant analyzer workflow, which reports the effect of a given set of variants (SNPs and small InDels) on genes and gene products without applying any mapping algorithm. This workflow supports read (FASTQ) and variant (VCF) files as input for the test sample and an optional control sample. The variant analyzer can be used to assess the effects of a short list of mutations as well as those identified in reads from whole-genome sequencing datasets. As in the previous workflows, the report includes tabular data and diagrams describing all variants contained within the input file. The following information is provided for each variant: its position in the genome, quality value (estimated by the variant-calling pipeline), read counts, allele frequency, nucleotide and amino acid changes (if present), gene and gene elements affected by the variant, functional annotation of the gene (if the corresponding functional annotation file is available), a pair of primer sequences that can be used to genotype the variant, and sequences flanking the variant in the reference genome.

#### Reporting of SNPs and small InDels

While the first version of Easymap only reported EMS-type mutations, the user can now specify if other type of SNPs or small InDels should be reported as candidates. This function allows these variants to be identified with the pre-existing linkage-analysis workflow, the newly implemented variant density mapping, QTL-seq mapping, and the variant analyzer workflows.

#### Flexibility of control samples

Easymap v.2 supports VCF files as control samples for all mapping analyses that do not require the computation of allele frequencies of the control sample variants. The use of VCF files instead of FASTQ files enables the use of a customized control sample consisting of a compilation of variants pooled from samples of different genotypes, as long as they are not linked to the phenotype of interest. This type of control sample is useful when working with strains with a high number of polymorphisms with the reference strain. The use of VCF files as control files also saves time for mapping analyses, since Easymap v.2 skips the time-consuming alignment and variant-calling steps for the control sample. Some mapping workflows implemented in Easymap v.2 can also be executed without a control sample. While this approach is highly unadvisable for most mapping scenarios, it can be useful for previsualizing data in the absence of a control sample.

#### Multi-threading

Easymap v.2 allows the user to set the number of dedicated central processing unit (CPU) threads for each analysis. This option is particularly useful when working with large genomes or large read files, as the analysis rate is proportional to the number of threads used during the steps that are compatible with multi-threading.

#### Preprocessing of reads

Preprocessing of NGS reads is a common step prior to any data analysis using FASTQ files. We incorporated the FASTQ preprocessing tool Fastp (Chen *et al.*, 2018) into Easymap v.2 as an optional step for every workflow, since it is fast, and easy to automate and implement within a bioinformatics pipeline. In Easymap v.2, fastp functions in its default configuration to perform automated quality filtering, adapter trimming, and read pruning, and can be enabled or disabled using a switch in the web interface prior to analysis.

## DISCUSSION

The increased availability and sharp decline in the cost of NGS technologies during the last decade has opened the door for researchers to use NGS on a semi-routine basis (Candela *et al.*, 2015; James *et al.*, 2013; Sarin *et al.*, 2008). However, manipulating NGS reads is a complex and time-consuming endeavor. Many tools and platforms have been developed for this purpose, but most are meant for bioinformaticians, as they require the user to combine multiple unrelated tools in order to perform a complete analysis. Specifically, for mutation mapping, few tools implement workflows that use raw reads to generate a list of candidate mutations in a user-friendly manner, and most of these tools lack versatility. The first version of Easymap was designed to ease mutation mapping by linkage analysis and to map large DNA insertions, making it quite useful for identifying transgenes and characterizing insertional lines of any type (Lup *et al.*, 2021). In Easymap v.2, we implemented additional workflows for other common mapping strategies.

Mapping approaches based on studying variant density in a pool of mutants that have been recurrently backcrossed to the reference strain are often used for *Caenorhabditis elegans* due to its short lifespan and the difficulty in isolating and phenotyping large subsets of individuals of the same generation (Svensk *et al.*, 2016; Zuryn *et al.*, 2010). These approaches are also used with large plants such as maize due to the spatial difficulty of simultaneously working with many individual plants (Klein *et al.*, 2018). We demonstrated the success of our variant density mapping workflow for datasets obtained using such approaches, especially when the test samples were in the reference background rather than in a highly divergent background. In the latter case, it is more difficult to discern between the causal mutation and non-causal variants regardless of the software used and, consequently, our variant density mapping workflow reported many candidates. This limitation can be addressed by using control samples that combine variants from multiple sub-samples into a VCF file, an option that is supported by Easymap v.2.

For QTL-seq mapping, automated workflows such as the one implemented in Easymap v.2 can rapidly point to genomic regions exhibiting linkage disequilibrium. Since QTL-seq relies on the use of two genetic backgrounds that are highly different from each other and from the reference genome, no control sample can be used to filter the data, as the variants of interest can be present in either of the two sequenced pools of individuals from the mapping population. Therefore, a vast number of candidate variants is commonly identified. Furthermore, the unknown molecular nature of the causal variants impedes any filtering step based on this property. In this sense, the identification of the causal gene is often disregarded in QTL-seq approaches due to the complexity of discerning between all the variants detected. Instead, narrow chromosomal regions are often defined, which can be used for the genetic improvement of crops (Fekih *et al.*, 2013; Illa-Berenguer *et al.*, 2015; Yang *et al.*, 2017). Further fine-mapping experiments such as linkage analysis to molecular markers and deep-sequencing are often required to narrow down the regions of interest or to identify the causal mutations, especially when working with large genomes and very low read depths (Wang *et al.*, 2020; Yang *et al.*, 2021).

Our software successfully identified the genomic regions harboring potential QTL in the tested datasets and reported all the variants, indicating those that could be of interest. Easymap v.2 provides lists of polymorphisms with detailed information to help users define narrower or alternative QTL-seq mapping intervals or to apply more stringent filters to detect candidate variants. Since the phenotype of interest could be caused by genetic variants that remain undetectable by re-aligning short reads to a reference genome, such as large InDels, microsatellites, or chromosomal re-arrangements (Doitsidou *et al.*, 2016), one list provided by Easymap v.2 contains the genes present in potential QTL to help the user identify additional candidates.

In conclusion, Easymap v.2 is a robust, versatile tool that can be used by researchers without previous experience in applying NGS strategies to gene mapping. Installing Easymap v.2 in any operating system is simple, as detailed in the one-page Quickstart Installation Guide (Supplementary File 1). Although the web interface is largely self-explanatory, comprehensive instructions and usage details can be found in the Easymap v.2 Documentation (Supplementary File 2). An interactive preview of the user interface with the mapping reports generated during the validation of all the workflows performed here is available at http://atlas.umh.es/easymapv2.

## DATA AVAILABILITY

Easymap is freely available at http://genetics.edu.umh.es/resources/easymap/. The sources of the datasets used in this work are detailed in Supplementary Tables 1 and 2.

## FUNDING

This work was supported by grants from the Ministerio de Ciencia e Innovación of Spain (PGC2018-093445-B-I00 and PID2021-127725NB-I00 [MCI/AEI/FEDER, UE]) and the Generalitat Valenciana (PROMETEO/2019/117) to JLM. SDL held a predoctoral fellowship (ACIF/2018/005) from the Generalitat Valenciana.

## Abbreviations

NGS: next-generation sequencing
EMS: ethyl methanesulfonate
SNP: single-nucleotide polymorphism
QTL: quantitative trait loci
VCF: Variant Call Format

## SUPPLEMENTARY DATA

SUPPLEMENTARY FILES

Supplementary File S1. Easymap v.2 documentation.

Supplementary File S2. Easymap v.2 quickstart installation guide.

SUPPLEMENTARY TABLES

Supplementary Table S1. Validation of Easymap v.2 for variant density mapping analyses.

Supplementary Table S2. Validation of Easymap v.2 for QTL-seq mapping analyses.

## Notes

### Competing Interest Statement

The authors have declared no competing interest.

