## Supplementary Files and Tables for "Versatile mapping-by-sequencing with Easymap v.2"

### Easymap v.2 Documentation

#### I. General description and scope

Easymap v.2 is an all-in-one software package that facilitates mutation mapping through the use of next-generation sequencing (NGS) reads in model organisms for which a reference genomic sequence is available (e.g., *Arabidopsis thaliana*, *Caenorhabditis elegans*…). It can be used from the command line or through a web graphical interface, in both cases either locally or remotely. Easymap v.2 is designed to map EMS-induced mutants carrying GC→AT transitions, natural variants detectable by short-read realigning to a reference genome (SNP and indels), and for mutants harboring large insertions such as transposable elements or T-DNAs. To map point mutations, Easymap v.2 uses genomic DNA or cDNA reads from a phenotyped $F_2$, $M_2$, or $M_3$ mapping population and from a control sample, and employs either linkage analysis or variant density mapping to identify one or several candidate regions. To map large insertions, it requires genomic DNA reads from one or several (pooled) insertional lines; mapping is performed by capturing genomic sequences flanking the insertions.

#### II. Availability and installation

Easymap v.2 is available under GNUv3 license for UNIX-based operating systems. For simple installation and environment configuration guidelines follow the Easymap Quickstart Installation Guide. For more detailed information, continue reading this documentation. For Windows users, Easymap v.2 can run in the Ubuntu app for Windows 10 available at the Microsoft Store. For macOS X users, Easymap v.2 can run in a virtual machine running Ubuntu. See section XIII for further details on different installation setups. Before installing Easymap v.2, check section X to determine whether other dependencies need to be installed beforehand. To install Easymap v.2 in a shared environment, see section XI. The following steps require administrator privileges.

Open a Linux terminal, go to the location in which to install Easymap v.2 (the `/home` directory in the example), download the source code, unzip it, and change permissions to allow installation (omit $ sign):

```
$ cd ~
$ wget
http://genetics.umh.es/other_files/genetica%20umh%20es/Easymap/easymap-v2.zip
$ unzip easymap-v2.zip
$ sudo chmod -R 755 easymap.v2
```

Now, to install the program, enter the `easymap.v2` directory and run the `install.sh` file.

Option 1 (to use Easymap v.2 exclusively through the command line):

```
$ cd easymap.v2
$ sudo ./install.sh cli
```

Option 2 (to use Easymap v.2 through the command line or its web interface):

```
$ cd easymap.v2
$ sudo ./install.sh server 8100
```

The second argument (8100) corresponds to the port in which the graphical interface of Easymap v.2 will be available. Use a port number between 8100 and 8200 that is not already in use; if no port number is specified, Easymap v.2 will by default use port 8100, which is typically available on most machines. Installation can take up to 30 minutes on a personal computer; on completion, a message indicates whether the installation was successful or not. If the installation was not successful, please review section X to make sure that all the required dependencies have been installed.

To uninstall the program, go to the `easymap.v2` directory and run the `uninstall.sh` script. After that, the whole `easymap.v2` directory can be manually removed.

```
$ cd /home/easymap.v2
$ sudo ./uninstall.sh
```

#### III. Input files required by Easymap v.2

File names must contain only alphanumeric characters and no blank spaces. All text files provided to Easymap v.2 should be formatted with UNIX line separators (`\n`); this is the case with FASTQ reads obtained from high-throughput sequencers or with FASTA and GFF3 files downloaded from biological databases (e.g., http://www.ensembl.org/info/data/ftp/index.html). However, if a file is edited in a Windows OS, it is worthwhile to check that the file is still readable by Easymap v.2 by opening it and inspecting it. If using the web interface, click on the "Preview" button that appears to the right of the file name (see section IV).

##### Mandatory input files

###### NGS data of the test sample in FASTQ format

If the reads are single end, provide one file (e.g., `sample.fastq`), whereas if they are paired end, provide two (e.g., `sample_1.fastq`, `sample_2.fastq`). FASTQ files have quality information associated with each nucleotide call; there are different encodings for this information, and Easymap v.2 needs the quality encoding to be "Sanger". Most NGS data produced in the past years have Sanger encoding characteristics. Nonetheless, Easymap v.2 inspects the input reads at the beginning of each execution and warns the user if their encoding is not Sanger. To check beforehand whether the reads meet this requirement, and to convert them if necessary, the easiest method is to analyze them with FastQC and convert them with FastQ groomer. Both tools can be found on the public Galaxy server (https://usegalaxy.org/).

For some mapping workflows Easymap v.2 needs two genomic DNA or cDNA read sets (test and control samples). It is possible to use different types of reads for the different samples (e.g., single-end reads for the test sample and paired-end reads from the control sample). Regarding read depth, Easymap v.2 will analyze datasets of any read depth, but values lower than 10× for insertional mutants and 25× for SNPs and small indels will compromise the accuracy of the results. Above these minimum values, the higher the read depth is, the more accurate and easier the interpretation of the results will be. The program reports the read depth distribution for each sample analyzed. Easymap v.2 also checks and reports the quality of the nucleotide calls in the provided reads. Be aware that low-quality calls can also compromise the results. If low-quality calls are at the 5′ or 3′ ends of the reads, consider trimming them with the appropriate software before performing an analysis with Easymap v.2.

###### Reference genome in FASTA format

If the reference genome has multiple contigs, its sequence can be provided as a single (e.g., `genome.fa`) or multiple FASTA files. In the second case, the file names must have the structure `{basename}.{contig_number}.fa` and share the same `base name` (e.g., `genome.1.fa`, `genome.2.fa`, etc). All the FASTA headers of the contigs must be present in the GFF3 file provided so that Easymap v.2 can link the information in the two files. FASTA and GFF files downloaded from the same databases are normally associated so that the names of the contigs coincide and no manipulation of the files is required. However, if the contig names in the FASTA and GFF files do not match, the FASTA files must be manually renamed. Easymap v.2 compares the input FASTA and GFF3 files at the beginning of each execution and warns the user if the headers in the FASTA file are not in the GFF3 file. Use all contigs of the reference genome even when knowing which one contains the mutation of interest; not doing so will increase alignment artefacts. For the same reason, include the reference sequences of organelles such as the mitochondrion (and chloroplast if applicable).

###### Gene structural annotation of the reference genome in GFF or GFF3 format

These files are available in the main biological sequences databases and are normally associated with reference FASTA files (http://www.ensembl.org/info/website/upload/gff3.html). If manipulating the GFF/GFF3 file, be sure not to not introduce additional characters such as " when saving the file, and be sure that the line separators are UNIX-like. This file is not mandatory for the QLT-seq mapping workflow in order to allow the use of non-annotated reference genomes. However, the mapping report and information provided by Easymap v.2 will lack any gene structural information (gene-altering mutations will not be reported as such).

##### Input files required for specific analyses

###### FASTA sequence of the insertion

This is only required for large-insertion mapping. The file must contain the full sequence of the insertion, but it can also contain additional sequences (e.g., the whole sequence of the vector used to engineer a transgene). If the file contains multiple FASTA headers, Easymap v.2 will use the first sequence.

**Optional input files**
**Control sample data in VCF format**
Control samples can be provided as a VCF (variant calling format) file. Easymap v.2 allows the use of VCF files of one sample or with variants pooled from different samples, however contig identifiers in the VCF file must match the contig identifiers in the genome reference file(s). Please check that this is the case and change the identifiers if needed. Read count and allele frequency information can be lost or missing from some VCF files, for this reason the variants will be only used to be filtered from the test sample, regardless of the allele frequency of the variants in the control sample. This impedes the use of VCF control files for the QTL-seq workflow and for linkage analysis mapping with a wild-type $F_2$ control sample.

**Gene functional annotation of the reference genome**
There is no standard format for this information, so Easymap v.2 asks for the simplest possible file: A tab-delimited text file with at least two columns, the first being the gene identifiers as found in the gene structural annotation file (e.g., At1g01010 for *Arabidopsis thaliana*), and the remainder containing their description (e.g., gene symbols, functional information, etc.). Except for the first one, columns can have blank records. The user can create tab-delimited text files with a spreadsheet editor by saving as “text (tab-delimited) (*.txt)”. Easymap v.2 can run without an annotation file, but it then will not include gene functional information in the mapping report. This file, or a very similar one that can be modified to fit the format specified above, is typically available from model organism databases.

### IV. Running Easymap v.2 through the web interface

#### Accessing Easymap v.2

If the machine has Easymap v.2 installed (local installation), point the web browser to http://localhost:<port-number> (e.g.: http://localhost:8100). If using Easymap v.2 remotely (hosted by institutional servers, in the cloud, or on a virtual machine), use the network IP where Easymap v.2 is installed: http://<ip-address>:<port-number> (e.g., http://11.22.33.44:8100). This address will be made available by the machine administrator. The port number is stored in the text file easymap.v2/config/port. Manual launch of the server might be required in exceptional cases. If at any time the interface cannot be accessed, use the following command to start the server manually:

```
$ cd /path/to/easymap.v2
$ bash launch-server.sh <port-number>
```

#### Uploading files to Easymap v.2

On the main menu, click on **Manage input files**, and then on **Select files**. This will open a browsing window of the operating system that allows the selection of multiple files at once. Easymap v.2 supports files with the following extensions: .fa, .fna, .fasta, .fq, .fastq, .gff, .gff3, and .txt. The program also takes gzip-compressed files with the extension .gz.

Click **Accept** and check that the files are listed ready for upload. Then, click **Upload files**. Easymap v.2 supports files of up to 200 Gb. After clicking on **Upload files**, the upload progress percentage of a file is shown; it increases up to 100%. After that, it may take a few minutes before the file appears listed and ready to use under the header “Current files in disk” on the **Manage input files** page. If uploading gzip-compressed files, it will take longer for the files to be ready because they must be unzipped.

Please do not close or refresh the **Manage input files** page during the file upload or de-compression, since this may interrupt the process. Upload speed to remote installations of Easymap v.2 depends on the network, and in exceptional cases, it might not be possible to upload files of several gigabases, even though Easymap v.2 features chunked file transfers for that purpose. In those cases, two alternatives to consider are using a local installation or transferring the files to the remote installation through the command line. For manual transfer of input files, they should be placed in the directory “/easymap.v2/user_data”.

#### Managing input files

On the main menu, click on **Manage input files** to see a list of the files currently uploaded to Easymap v.2. To preview a file, click on the **Preview** button next to its name. This will open a new browser tab with the first 1000 lines of the file. This is useful when checking the content of very large files (e.g., FASTQ read files) that most machines are unable to open. To delete a file from the easymap.v2 directory click on the button **Remove from disk** next to its name. This will display a warning message requesting confirmation. Click on **Confirm** to delete the file permanently.

#### Running a project

A project is an Easymap v.2 execution using a given set of files and performing particular tasks based on the user’s preferences. On the main menu, click on **Run new project** and choose the preferred files and mapping options. Then, click on **Check input and run project**. This will check the user input and, if valid, will unlock the button **Start project**. Review all the options and click on that button when ready. This will launch the project and redirect the user to the **Manage projects** menu, where there is a new project running. All fields on screen are mandatory except **Gene functional annotation file**. The options are largely self-explanatory, assistance is provided on-screen, and Easymap v.2 will check all the choices made by the user. However, for reference, here is a description of each field available:

1. **Project name**: Provide a meaningful name to identify the project. Blank spaces and non-alphanumeric characters are not allowed as this name will be used to create a directory to store the files related to this project. Easymap v.2 will automatically append a timestamp to the project name to make it unique. Use this name to monitor and review the project in the **Manage projects** menu.
2. **Mapping-by-sequencing strategy**: Choose between **Linkage analysis mapping**, **Variant density mapping** or **QTL-seq** for point mutation mapping, **Tagged sequence mapping** for large insertion mapping or **Variant analyzer** to analyze the effect of all variants in a sample without applying a mapping algorithm.
3. **Data source**: Only available for **Tagged sequence mapping** and **Linkage analysis mapping**. Choose between **Use my own reads**, to analyze experimental reads obtained in a high-throughput sequencer, or **Simulate data**, for Easymap v.2 to simulate reads (see section XII).

4. **Reference sequence**: Choose the base name of the reference genome to use as template. Easymap v.2 will then automatically select all the files that have the same base name (see section III). If the reference files are not listed in the dropdown menu, check that their names have a "`.fa`", "`.fna`" or "`.fasta`" extension.
5. **Insertion sequence file**: Only available for **Tagged sequence mapping**. Choose the FASTA file that contains the sequence of the insertion of interest. If this file is not listed in the dropdown menu, check that its name has a "`.fa`" extension.
6. **GFF3 file**: Choose the GFF3 file that contains the structural annotation of the reference genome. If this file is not listed in the dropdown menu, check that its name has a "`.gff`" or "`.gff3`" extension. For **QTL-seq** strategies, the user may select "None" if no structural annotation file is available for the reference genome, however this restricts the information Easymap v.2 can report.
7. **Gene functional annotation file**: This field is optional. Choose a file that contains functional information about the genes in the GFF3 file. If this file is not listed in the dropdown menu, check that its name has "`.txt`" extension.
8. **Mutant background**: Only available for **Linkage analysis mapping**. Choose between **Reference** and **Non-reference**. Pick the former when the genetic background of the mutant under study is the same as the sequence provided in **Reference sequence**, and otherwise pick the latter. Only certain combinations of **Mutant background**, **Mapping cross performed**, and **Origin of the control reads** are allowed (see Table 1).
9. **Mapping cross performed**: Only available for **Linkage analysis mapping**. Select between **Backcross** and **Outcross**. Choose the former if the mapping population was obtained by crossing the mutant with its premutagenesis parental (or equivalent), and the latter if it was crossed with a strain polymorphic to its premutagenesis parental.
10. **Origin of the control reads**: Only available for **Linkage analysis mapping**. Select between **Mutant parental**, **Polymorphic strain**, and **$F_2$ wild types**. "Mutant parental" is the strain mutagenized with EMS, "Polymorphic strain" is the strain crossed with the mutant in an outcross, and "$F_2$ wild types" is a bulked group of $M_2$ or $F_2$ plants that do not show the recessive phenotype. Select the sample to be used as a control during the analyses. See Table 1 to know which samples can be used as control for a particular mapping experimental design in Easymap v.2.
11. **Use low stringency during SNP analysis?**: Available for **Linkage analysis mapping**, **Variant density mapping** and **QTL-seq**. By default, Easymap v.2 considers only SNPs that pass certain quality checks. However, with some non-optimal datasets, more lenient filtering can help identify a candidate interval and the causal mutation. If the analysis was performed in the default mode and rendered very few SNPs and small indels or the causal mutation might have been discarded by the program, turn on this option and run the program again.
12. **Test data**: Choose the file(s) that constitutes the test sample. If the reads are single end, select only one file; if they are paired end, select two files while holding down the Ctrl/Cmd key. For the **Variant analyzer** workflow, VCF files will be supported as well.
13. **Control data**: Choose the file(s) that constitutes the control sample. For some cases, the test sample can be provided in VCF format, otherwise FASTQ read files will be requested. If the reads are single end, select only one file; if they are paired end, select two files while holding down the Ctrl/Cmd key. For the **Linkage analysis mapping** workflow, these files must correspond to the control sample specified in **Origin of the control reads**. For the **Variant density mapping** workflow, the variants in this sample will be subtracted from the test sample. For the **QTL-seq** workflow, FASTQ reads of one of the sequenced populations should be selected.
14. **Mutagenesis simulation parameters**: Only available when selecting **Simulate reads**. This field consists in a preformatted JSON string (`{"numberMutations":"0"}`). Replace the values with the desired ones. See section XII for more information.
15. **Recombination and selection simulation parameters**: Only available when selecting **Simulate reads** and **Linkage analysis mapping**. This field consists in a preformatted JSON string. Replace the values with the desired ones. See section XII for more information.
16. **Contig recombination frequency distributions**: Only available when selecting **Simulate reads** and **Linkage analysis mapping**. This field consists in a preformatted JSON string. Replace the values with the desired ones. See section XII for more information.
17. **Sequencing simulation parameters**: Only available when selecting **Simulate reads**. This field consists in a preformatted JSON string. Replace the values with the desired ones. See section XII for more information.

If the maximum space allowed for Easymap v.2 or the maximum number of simultaneous jobs running have been surpassed, a red warning banner will appear at the top of the page, and the ability to run new projects will be temporarily disabled. This is a safety limit imposed by the administrator of the machine and can be configured (see section XI).

**Managing projects**

On the main menu, click on **Manage projects** to view all the projects stored in the disk from the most recent to the oldest. The status of every project, and the amount of memory it uses, are displayed. Project statuses can be “running”, “finished” (all tasks finished without errors), “killed” (the user stopped the project before completion), or “error” (one or more tasks returned an error). The options available for a project depend on its status and are displayed as buttons. Here is a description of these options:

**View log file**: Points the browser to a live log file. If a project is running, it can be useful to know what tasks are already completed. If a project has finished, this can be used to review what tasks were performed and how long each one took. If a project returns an error, the log file shows an explanation of the error and in which task it occurred. An execution can require tens or hundreds of gigabases during certain steps. If the log of an uncompleted project indicates that it is halted, check that there is enough free space on the disk.
**Stop project**: Aborts a running project. A warning message asking for confirmation appears. Click on **Confirm** to stop the project irreversibly.
**Remove from disk**: Deletes all files generated by a project. This button is only available if the project is not running. To delete a running project, stop it first. A warning message asking for confirmation appears. Click on **Confirm** to delete the project irreversibly.
**View report**: Points the browser to the project report. This button is only available after all the project tasks have been completed. See section VIII for a detailed explanation of the mapping reports.

#### V. Quick start reference to run Easymap v.2 through the command line

Place all the input files in `easymap.v2/user_data`. All files must be unzipped. To specify the input files in the Easymap v.2 command, simply type the name of the file, not the path to it. For more information about the required input files, see section III. Easymap v.2 can map any SNP, small indels, and large insertions. Section VI contains a table that describes all the arguments for the Easymap v.2 command, but for a quick start, continue reading here.

To map insertions with single-end reads, use this command:

```
./easymap -w ins -n <string> -r <string> -i <file> -p <file> -g <file> -a <file>
```

where `-w ins` selects a workflow to map insertions, `-n` indicates the project name, `-i` the insertion file, `-r` the reference genome basename, `-p` the test reads, `-g` the GFF3 file, and `-a` the annotation file. The `-a` argument is optional, so in the absence of a functional annotation file, simply omit it. Example:

```
./easymap -w ins -n myProject -r celegans -i Mos1 -p mutReads.fq -g ceGenes.gff -a ceAnnotation.txt
```

If the reads are paired end, specify the names of both files separated by a comma:

```
./easymap -w ins -n myProject -r celegans -i Mos1 -p mutReadsF.fq,mutReadsR.fq -g ceGenes.gff -a ceAnnotation.txt
```

In the following example of a linkage analysis mapping, (a) an EMS-induced mutant in the reference background (b) was backcrossed to its wild-type parent to create a mapping population, (c) from which a pool of phenotypically mutant $F_2$ plants was sequenced, (d) and reads from the mutant parent were used as control. The mapping is performed with the following command:

```
./easymap -w snp -n myProject -r athaliana -g atGenes.gff -a atAnnotation.txt -p F2mutF.fq,F2mutR.fq -c wtRefF.fq,wtRefR.fq -ed ref_bc_parmut
```

where `-c` specifies the control reads and `-ed` the experimental design that was used to obtain the reads. For the list of experimental designs to map SNPs and small indels supported by Easymap v.2, see section VII.

Easymap v.2 comes with a simulator module that can simulate linkage analysis mapping and tagged sequence mapping scenarios. As an example, to simulate and analyze reads for a virtual mutant with one randomly positioned insertion, edit the file `/easymap.v2/simulator/sim_parameters.json` to specify the simulation parameters:

```
{
    "SimMut": {
        "NumberMutations":"1"
    },
    "SimRecsel": {
        "ContigCausalMutation":"1",
        "PositionCausalMutation":"30000",
        "NumberRecombinantChromosomes":"100",
        "RecombinationFrequencies": {
            "1":"0,25-1,32-2,34-3,15-4,5-5,2",
            "2":"0,25-1,40-2,23-3,5-4,1-5,1",
            "3":"0,21-1,40-2,29-3,10-4,3-5,1"
        }
    },
    "SimSeq": {
        "Library":"pe",
        "FragmentSize":"500",
        "FragmentSd":"100",
        "ReadDepth":"20",
        "ReadSize":"100",
        "ReadSd":"0",
        "ErrorRate":"1",
        "GCbias":"50"
    }
}
```

`"NumberMutations":"1"` creates one insertion in the reference genome (see section XII for an explanation of each JSON “`name`”:“`value`” pair). Save the file and run the following command:

```
./easymap -w ins -n myProject -r celegans -i Mos1.fa -g ceGenes.gff -a ceAnnotation.txt -sim
```

The `-sim` flag turns on the Easymap v.2 built-in simulator. Note that now there is no `-p` (test reads) argument.

During the execution of Easymap v.2, all output is redirected to `easymap.v2/user_projects/{timestamp}_project_name/2_logs/log.log`. This file and other log files in the same directory can be checked to monitor the execution of a project. An execution can require tens or hundreds of gigabytes during certain steps. If the log indicates that the project halted, check that there is enough free space in the disk. If the project halts returning an error, other files in addition to the log can be checked under `easymap.v2/user_projects/{timestamp}_project_name/2_logs/`. Once an Easymap v.2 job has finished successfully, review the results in `easymap.v2/user_projects/{timestamp}_project_name/3_workflow_output/`. See section VIII for a complete description of the Easymap v.2 report and all the output datasets it produces.

#### VI. Full list of Easymap v.2 command line arguments

| | |
|---|---|
| `-w, --workflow`<br>Type: string (`snp`, `ins`)<br>Required: always | Analysis workflow. Use `snp` for linkage analysis mapping, `ins` for tagged sequence mapping, `dens` for variant density mapping, `qtl` for QTL-seq mapping and `vars` to use Easymap v.2 as a variant analyzer. |
| `-n, --project-name`<br>Type: string<br>Required: always | String containing the name of the project. Only alphanumeric characters are allowed. The files of the project are located in `easymap.v2/user_projects/{timestamp}_project_name`. |
| `-r, --reference-sequence`<br>Type: string<br>Required: always | Base name of the FASTA file or files that contain the reference sequence. Regardless of whether there are one (e.g., `danio_rerio.fa`) or multiple files (e.g., `danio_rerio.1.fa`, `danio_rerio.2.fa`, …), type `danio_rerio`. |
| `-i, --insertion-sequence`<br>Type: file<br>Required: only if `-w ins` | Name of the FASTA file that contains the insertion sequence. It supports files with numbered lines and/or blank lines. |
| `-g, --gff3-file`<br>Type: file<br>Required: always except when `-w qtl` | Name of the GFF3 file that contains the genome structural annotation. It is always required, except for QTL-seq mapping, where this argument is optional. The contig identifiers in this file must match the contig identifiers in the genome reference file(s). |
| `-a, --annotation-file`<br>Type: file<br>Required: optional | Name of the file that contains functional information of the genes in the reference genome. The format is custom (see section III). |
| `-p, --reads-test`<br>Type: file<br>Required: when `-sim` is off | Name of the FASTQ file(s) containing the reads of the test sample. If the reads are paired end, supply both file names separated by a comma without spaces (`readsF.fq,readsR.fq`). When `-w vars`, the workflow supports either a FASTQ or VCF file. |
| `-c, --reads-control`<br>Type: file<br>Required: optional | Name of the FASTQ or VCF file(s) that constitutes the control sample. See section VII to know which files can be used as control sample. If the reads are paired end, supply both file names separated by a comma without spaces. |
| `-ed, --experimental-design`<br>Type: string<br>Required: only if `-w snp` | For `-w snp`, string that summarizes the experimental design followed to obtain the read datasets provided to Easymap v.2. See Table 1 to know which string matches the experimental design. |
| `-emt, --exp-mut-type`<br>Type: string | To specify whether only EMS-type mutations or all detected mutations should be reported when `-w snp`, `-w qtl`, `-w vdm`, or `-w vars`. Use “EMS” for EMS-type mutations and “all” for all detected mutations. Default = “EMS”. |
| `-ls, --low-stringency`<br>Type: flag | By default, Easymap v.2 considers only SNPs that pass certain quality checks. However, in some read datasets that are not optimal, performing more lenient filtering can help to identify a candidate interval and the causal mutation. If the read analysis in the default mode renders very few SNPs or the causal mutation could have been discarded, turn on this option and run the program again. |
| `-tr, --threads`<br>Type: integer<br>Required: optional | Number of CPU threads to use for workflow steps compatible with multithreading. Default = 1. |
| `-sim, --simulate-reads`<br>Type: flag | Only with `-w snp` and if `-w ins`. Turns on Easymap v.2 built-in function to simulate the data for a mapping experiment. See section XII for details of how to specify the simulation parameters. |
| `-ppc, --preprocessing`<br>Type: flag | Activates preprocessing of input reads with a default run of “fastp”. |
| `-u, --usage`<br>Type: flag | Shows where to find help to run Easymap v.2 through the command line interface. |

#### VII. Experimental designs supported by Easymap v.2 for linkage analysis mapping of point mutations and small indels

Easymap v.2 requires two read datasets for mapping point mutations. The first corresponds to a pool of phenotypically mutant individuals from an $M_2$ generation or from the $F_2$ generation of a mapping cross. The second corresponds to a control sample and depends on the background of the mutant under study and the source of the mapping population. Since Easymap v.2 includes a splice-site aware aligner, both datasets can come from a DNA-seq or an RNA-seq sample. See Table 1 to know which samples can be used as control in each case.

**Table 1.-** Experimental designs for linkage analysis mapping of SNPs and small indels supported by Easymap.v2

| Genetic background of the mutant | Source of the mapping population | Test sample | Control sample | Argument value `-ed` / `--experimental-design` |
|---|---|---|---|---|
| The same as the reference sequence provided | $M_2$ or backcross $F_2$ | DNA or RNA from a pool of $M_2$/$F_2$ phenotypically mutant individuals | Mutant parent | `ref_bc_parmut` |
| | | | Pool of $M_2$/$F_2$ phenotypically wild-type individuals | `ref_bc_f2wt` |
| | Outcross $F_2$ | | Mutant parent | `ref_oc_parmut` |
| | | | Polymorphic strain | `ref_oc_polstr` |
| Other than the reference sequence provided | $M_2$ or backcross $F_2$ | | Pool of $M_2$/$F_2$ phenotypically wild-type individuals | `noref_bc_f2wt` |
| | Outcross $F_2$ | | Mutant parent | `noref_bc_parmut` |

##### VII.1. SNP selection and filtering parameters

**Linkage analysis mapping**

Easymap v.2 selects high-confidence polymorphisms for mutation mapping according to the following criteria: (1) SNPs with quality values as defined by the variant-calling pipeline over 100 for the high stringency mode and over 30 for the low stringency mode. (2) SNPs with a reasonable read depth, avoiding excessively low or high values as compared with the overall average read depth. For datasets with an average read depth over 25, Easymap v.2 filters out SNPs with an RD lower than 15. For datasets with an average RD below 25, Easymap v.2 filters out SNPs with an RD below 10. For datasets with an average RD over 40, Easymap v.2 filters out SNPs with an RD greater than three times the average RD, while for datasets with an RD under 40, Easymap v.2 filters out SNPs with an average RD over 100. (3) EMS-type mutations for mutants in a reference genomic background backcrossed with the mutant parental line. These arbitrary thresholds have proven to be effective in selecting high-confidence polymorphisms for mutation mapping, avoiding conflicting SNPs.

For candidate mutation reporting, Easymap v.2 applies the following filters: (1) SNPs exclusive to the test sample, (2) contained within the candidate region, (3) with an allele frequency higher than 0.8 and (4) EMS-compatible mutations if this option is selected.

**Variant density mapping**

This workflow produces a sequence of sub-filtered lists of variants: total variants in the test sample; variants specific to the test sample (filtered with a control sample); EMS-type test sample specific variants; homozygous test sample specific variants; and EMS-type homozygous test sample specific variants. General filtering parameters for all sub-lists are: (1) quality values as defined by the variant-calling pipeline over 100 for the high stringency mode and over 30 for the low stringency mode; (2) RD over 12 for datasets with an average RD over 25, RD over 6 for datasets with an average RD under 25; (3) for datasets with an average RD over 40, Easymap v.2 filters out SNPs with an RD greater than three times the average RD, while for datasets with an RD under 40, it filters out SNPs with an average RD over 100.

For candidate mutation reporting, Easymap v.2 applies the following filters: (1) SNPs exclusive to the test sample, (2) contained within the selected regions, (3) with an allele frequency higher than 0.7 and (4) EMS-compatible mutations if this option is selected.

**QTL-seq mapping**

Easymap v.2 selects SNPs for mapping according to the following criteria: (1) quality value (as defined by the variant-calling pipeline) over 100 in the test sample, (2) present in the test and control samples, (3) read depth over 10 in both samples, (4) allele frequency cannot be 1 in both samples (this workflow eliminates non-segregating SNPs), and (5) allele frequency above 0.2 in both samples. Easymap v.2 reports mutations with a |dAF| > 0.4 that affect coding sequences. This workflow also reports a list of all genes contained within the selected regions.

**Variant analyzer**

This workflow applies no filters to VCF input data, since it generally consists in a list of polymorphisms from which to study their effects on genes. For FASTQ input files, minimal filters are applied, selecting variants with a quality value over 30 and a read depth over 4.

#### VII.2. Mapping algorithms general overview

**Linkage analysis mapping**

Easymap v.2 identifies candidate regions from different sources of data belonging to different experimental designs. First, the pipeline divides the genome into overlapping windows, the size and degree of the overlap varies according to the experimental design. Next, the SNP allele frequencies are averaged in each window and transformed in a Boost value if the experimental design includes an outcross. Average allele frequencies or Boost values are then revised and corrected by comparing up to 6 adjacent windows to reduce noise and randomly generated peaks using a weighted average approach. Easymap v.2 analyzes the revised allele frequencies or Boost values in each window to select the one with the highest value as the most likely to contain the candidate mutation, which will define the phenotypically selected genomic position or center of the candidate interval. Finally, a candidate interval of 4, 10, or 20 Mb is defined according to the size of the input genome, and the mutations within the candidate interval are reported as candidates.

**Tagged sequence mapping**

Easymap v.2 uses two complementary algorithms for large insertion mapping: (1) local alignment analysis relies on the selection of reads that can be partially aligned to the sequence of the DNA insertion (local alignments); (2) paired-read analysis is only used when paired-end reads are provided and relies on the selection of reads whose mate read can be aligned to the sequence of the DNA insertion. Then, in both algorithms, the reads are also aligned to the reference genome, generating clusters of reads around the position of each insertion. For a cluster to be considered a true positive and reported as an insertion site it must fulfill at least two of the following criteria: (1) have forward and reverse supporting reads, (2) have at least one position of the cluster with a RD >= 3, (3) cover a span of at least 200 bp.

**Variant density mapping**

This workflow relies on the selection of genomic positions with a density of variants higher than the rest of the genome. The genome is divided into 1-Mb windows, and the number of variants in each window is counted. These values are then revised and corrected by comparing up to 4 adjacent windows to reduce noise and randomly generated peaks using a weighted average approach. Easymap v.2 analyzes the revised values to select the highest value in each list of variants, which is then used to define additional peaks across the genome by selecting windows with a number of variants that is higher than 0.8 times the highest value. Once a list of peaks is defined, left and right limits are defined for each peak taking into account its adjacent windows.

**QTL-seq mapping**

SNPs that are informative for mapping are selected from both samples and the difference between allele frequencies in each sample is computed (dAF). dAF values are then averaged in 100-kb windows across the genome and mean dAF values are corrected with weighted average taking into account up to 4 adjacent windows. Corrected values are then used to select genomic regions with linkage disequilibrium, arbitrarily defined as regions with a corrected |dAF| > 0.4. If no region is selected, the threshold is lowered by 0.05 until a region is selected or until a threshold of 0.2 is reached.

#### VIII. Explanation of the Easymap v.2 output report

The Easymap v.2 output report consists in an HTML document, independently of whether Easymap v.2 was executed through the command line or through the web interface. To view the report of a project generated through the command line, go to `easymap.v2/user_projects/{timestamp}_project_name/3_workflow_output/report.html`. To view it through the web interface, go to **Manage projects**, locate the desired project, and click on **View report**. The report file is only available when the project status is "finished". To create a PDF or paper copy of the report, use the printing menu of the web browser, which will print an optimized version of the report. The report contains all the relevant information of the project, and it is organized in different sections. The main sections, which are common to most of the workflows, are the following:

1. **Run summary**: This section contains general information about the project, including the name of the input files and the options selected for the mapping analysis.

2. **Input data quality assessment**: Easymap v.2 performs a few checks on the quality of the provided data to assess the reliability of the mapping results:
   **Test and control sample read depth distributions**: A plot with the distribution of read depth frequencies found during the alignment of the reads to the reference genome gives a visual estimation of the average read depth in the test and control samples. Samples with low average read depth render less reliable mapping results.
   **Reads quality assessment**: A box-plot representing the variation of the quality of the reads along the read length. Low-quality reads hinder the mapping analysis, except in the case of the variant analyzer workflow, which does not perform any mapping analysis.

3. **Mapping or density analysis overview**: This section displays all input contigs along with the polymorphisms used for mapping, and the regions of interest are highlighted. The display is adapted to each workflow. This section is absent in the Variant analyzer report.

4. **Candidate region or variant analysis**: This section consists in a table that summarizes the information about the candidate polymorphisms or insertions. Each row corresponds to one polymorphism and contains the following information:
   **ID**: Numeric identifier assigned by Easymap v.2 to each candidate. It can be used to correlate information within the mapping report.
   **Contig, Position**: The contig and the absolute position of the polymorphism.
   **AF**: Frequency of the non-reference allele in the test sample.
   **Distance to peak (DTP)**: Distance between the polymorphism and the selected genomic position determined by Easymap v.2. Polymorphisms to the left and right sides of the selected position have negative and positive DTP values, respectively.
   **Nucleotide (Ref/Alt)**: The reference and alternative alleles of the polymorphism.
   **Gene (gene element)**: Gene identifier, as given in the input GFF file, and the element of the gene that contains the polymorphism (coding sequence [cds], intron, promoter, etc.). Except in the Tagged sequenced mapping report, Easymap v.2 reports putative splicing signal modifications located 2 nucleotides in the intron borders and 1 nucleotide in the exon border. Other intronic regions, such as branching sequences, are not analyzed.
   **Amino acid (Ref/Alt)**: When a polymorphism results in a change in a protein sequence, the reference and alternative sequences are given.

Additional information can be seen by clicking on the links provided. Those links open a text file containing all the available information regarding the candidate polymorphisms or all the polymorphisms in the test sample. These files contain the following additional fields:

**quality**: Phred-scaled quality score for the polymorphism as given by the variant calling pipeline.
**ref_count**: Number of reads containing the reference allele of the variant.
**alt_count**: Number of reads containing the alternative allele of the variant.
**hit**: Indicates if the mutation is located within a transcription unit (tu), in a regulatory region (rr), or in intergenic regions (nh, no hit).
**mrna_start, mrna_end**: Genomic coordinates of the transcription unit containing the mutation.
**strand**: Indicates the orientation of the gene within the contig.
**gene_funct_annot**: Gene functional annotation. If a functional annotation file is provided, this column contains information regarding the gene that contains the variant.

**f_primer, r_primer**: Forward and reverse primers designed for genotyping purposes. The amplified fragment should have a length of around 800 nt, with the polymorphism at a distance of approximately 300 nt from the forward primer.
**tm_f_primer, tm_r_primer**: Melting temperatures of the forward and reverse primers. Melting temperatures range between 60 and 64°C.
**upstream, downstream**: 50 nucleotides of DNA sequence upstream and downstream of the position of the variant.

5. **Candidate variants or Detected insertions**: The “Candidate variants” section is common to the Linkage analysis mapping, Variant density mapping, QTL-seq mapping, and the Variant analyzer workflows. This section contains a list of the candidate mutations affecting gene open reading frames. An image is generated representing each candidate gene: exons and untranslated regions are shown as boxes (dark and light blue respectively) connected by introns shown as lines. A 250-nt putative promoter region is included as a dashed line. The position of the variant is indicated with a red arrow accompanied by the nucleotide change and the amino acid change (when applicable). Finally, the image contains a scale bar to be used as a visual reference. The image is followed by a table containing relevant information about the variant and the affected gene. The “Detected insertions” section is found in the Tagged sequence mapping report, and it is an expanded version of the “Candidate variants” section (see section VIII.2 for further information).

Although the “Run summary” and the “Input data quality assessment” sections are identical in all the reports, the other sections contain specific data according to the analysis performed. The next paragraphs detail the peculiarities of the sections contained in each report.

#### VIII.1. Linkage analysis mapping report

**Section 3: Mapping analysis overview**

In this section, all input contigs are displayed along with the polymorphisms used for mapping the causal mutation and the linear descriptor of the allele frequency. The plot shows the position of each polymorphism in the contig (*x* axis) and the allele frequency of the non-reference allele (*y* axis). The candidate region determined by the mapping analysis is highlighted in pink, and polymorphisms from the test sample and the control sample are drawn in blue and orange, respectively.

**Section 4: Candidate polymorphisms overview**

The contig containing the candidate region is displayed along with all of the polymorphisms in the test sample that pass the quality checks. The window that contains the candidate mutations is highlighted, and the selected chromosomic position is represented as a dashed line. This window only includes polymorphisms with an allele frequency higher than 0.8. An additional image zooms into the candidate window displaying the final candidate polymorphisms.

**Section 5: Candidate region analysis**

This section summarizes the information regarding the candidate polymorphisms. Additional information can be seen by clicking on the “extended information” or the “all variants” links, located below the table.

#### VIII.2. Tagged sequence mapping report

**Section 3: Mapping analysis overview**

**Subsection 3.1: Genomic overview**

A representation of the input contigs with the positions of all detected insertions marked with red triangles.

**Subsection 3.2: Insertions summary**

The elements contained in this table are:

**Ins:** Numeric identifier given to each insertion.
**Contig, Position:** The contig and the absolute position of the insertion..
**Gene (gene element):** Gene name as given in the input GFF file, and the element of the gene that is interrupted by the insertion (coding sequence [cds], intron, promoter, etc.).
**Wt amino acids:** Number of wild-type amino acids conserved in the mutant protein.

Additional information can be seen by clicking on the "extended information" link, which opens a text file with all the available information regarding the insertions. This file contains the following fields, in addition to those already explained in section VIII):

**5_end_ins, 3_end_ins**: Reconstructed sequences for the 5′ and 3′ ends of the insertion as found in the mutant genome. The sequences are used to generate primers for genotyping the insertion.

**insertion_primer_5, insertion_primer_3**: Primers contained within the 5′ and 3′ ends flanking the insertion, and designed for genotyping purposes.

**tm_insertion_primer_5, tm_insertion_primer_3**: Melting temperatures of the described primers. Melting temperatures range between 60 and 64°C.

**Section 4: Detected insertions**

This section contains detailed information about each insertion, starting with histograms that visually summarize the mapping information. The histograms plot the read depth versus genomic position of the reads that support the detection of each insertion. When paired-end reads are available, two histograms are shown: (1) Flanking unpaired alignments: Reads that cannot be aligned to the insertion sequence, but whose complementary reads can; they surround the position of the insertion site in the genome. (2) Flanking local alignments: Reads that are aligned locally to the insertion sequence are then realigned to the genome sequence, marking the position at which the insertion event has occurred. When single-end reads are used, only the Flanking local alignments histogram will be available.

If a given insertion interrupts one or more genes, the information about each one is listed below the histograms, starting with a graphical representation of the gene in which the location of the insertion is marked with a red triangle. The image is followed by relevant information about the insertion, such as its position, the functional annotation (when available), the genotyping primers, and the genome sequences flanking the insertion site.

#### VIII.3. Variant density mapping report

**Section 3: Density analysis overview**

A representation of the input contigs plotting the number of variants per 1-Mb window in different filtered sub-lists of variants. Regions of interest are highlighted.

**Section 4: Candidate region analysis**

This section contains a table in which each row is a candidate variant with an allele frequency over 0.7 in the test sample and is absent from the control sample. Additional information can be seen by clicking on the "a list of all variants in the selected region" or "all variants in the genome" links, located below the table.

#### VIII.4. QTL-seq mapping report

**Section 3: Mapping analysis**

All input contigs are displayed, polymorphisms used for mapping are represented as black dots using their dAF and genomic position as coordinates in each plot. Corrected average dAF values are represented as a red line, and regions showing linkage disequilibrium are highlighted in pink.

**Section 4: Candidate variants overview**

This section displays a table in which each row is a candidate variant affecting a coding sequence with a |dAF| over 0.4. Instead of the field **AF** found in the reports of other workflows (see section VIII), this table contains the following:

**AF_test, AF_control**: Frequency of the non-reference alleles in the test and control samples respectively.

**dAF:** AF_test – AF_control.

Additional information can be seen by clicking on the "all variants within the highlighted QTL" or "all variants in the genome" links, found above the table. Furthermore, the link "a list of all genes within the highlighted QTL" provides a list of all genes contained within the selected regions.

#### IX. Content of the project directory

Although the output report contains all the relevant information of the project, additional files such as individual image files and tab-delimited text files can be accessed in the project directory through the web browser, the command line, or the operating system file manager. The project directory contains the following three folders:

**1_intermediate_files**: Contains intermediate files generated during the execution of a mapping analysis. Raw data and large intermediate files, such as SAM, BAM, or VCF files, are automatically deleted during the execution of the program to save storage space.

**2_logs**: This folder stores log files automatically produced by third-party software such as bowtie2 and SAMtools, as well as the Easymap v.2 log file for the given project (log.log).

**3_workflow_output**: Contains the report file (report.html), the images shown in the report, and additional images that might be of interest. This folder also contains the tab-delimited text files referenced in the report file (`candidate_variants.txt`, `candidate_variants_total.txt`). The names of all files in this folder are self-explanatory.

#### X. Appendix A: Dependencies needed to run Easymap v.2

All third-party software needed for the analyses is included in Easymap v.2; however, a few basic Linux packages are required for the installation of Easymap v.2. To prepare the system for the installation, please follow these steps.

In Ubuntu 20, run the following commands:

```
$ sudo apt-get update
$ sudo apt-get install build-essential zlib1g-dev libbz2-dev git wget tar zip liblzma-dev libncurses5-dev libncursesw5-dev libssl-dev make -y
```

In Ubuntu 18, run the following commands:

```
$ sudo apt-get update
$ sudo apt-get install build-essential zlib1g-dev libbz2-dev liblzma-dev libncurses5-dev libncursesw5-dev libssl1.0-dev wget tar zip git
```

In older versions of Ubuntu (tested in 16.04 and 14.04), run the following commands:

```
$ sudo apt-get update
$ sudo apt-get install build-essential zlib1g-dev libbz2-dev liblzma-dev libncurses5-dev libncursesw5-dev libssl-dev wget tar zip git
```

In RPM-based Linux distributions (Tested in Red Hat) run the following commands:

```
$ sudo yum groupinstall "Development Tools"
$ sudo yum groupinstall "Development Libraries"
$ sudo yum install wget zlib-devel bzip2-devel ncurses-devel ncurses openssl-devel
$ sudo yum install -y xz-devel
$ sudo yum install curl
$ sudo yum install curl-devel
```

#### XI. Appendix B: Installing Easymap v.2 in a shared environment

The installation script sets unrestricted permissions recursively in the `easymap.v2` directory. Change them manually after the installation if more restrictive permissions are needed. When installing Easymap v.2 for other users, the administrator can limit the resources available to them by editing the file `easymap.v2/config/config`. Further instructions can be found inside the file.

The installation script installs `python2.7` locally under the `easymap.v2` root directory to make sure it is available for Easymap v.2 without interfering with other installations that might already be on the machine. It also installs Virtualenv locally for the same reason. After that, it creates a virtual environment named `easymap-env` to isolate any other python libraries such as Pillow.

To make the Easymap v.2 web interface easily available for most users, the installation command `./install.sh server <port-number>` starts the python CGIHTTPServer in the background and modifies `/etc/crontab` so that the server is also started after each reboot. To avoid that, use `./install.sh cli`. Easymap v.2 includes an additional web interface in `easymap.v2/web_interface_PHP` based on a HTML-Javascript-PHP stack easily deployable in servers such as Apache2 that have PHP enabled.

#### XII. Appendix C: How to simulate data with Easymap v.2

To simulate data through the command line, include `--simulate-data/-sim` in the command and specify the simulation parameters by editing the file `easymap.v2/simulator/ sim_parameters.json`. The fields included in the file can also be edited through the graphical interface. Do not change the name of the file. A simulation of an insertional mutant consists in the creation of a mutant FASTA and high-throughput FASTQ reads. A simulation of an EMS mapping population and a control sample also involves the creation of recombinant FASTA files.

| | |
|---|---|
| `"NumberMutations"` | Specifies the number of mutations in the mutant. The reference sequence provided with `-r` is used as template to create a mutant version. If `–w ins`, the insertion sequence provided in `-i` is used as insert, and the number of insertions is limited to 1 per megabase of reference genome. If `–w snp`, the number of point mutations is limited to 100 per megabase of reference genome. Both insertions and point mutations are generated at random positions. |
| `"ContigCausalMutation"`<br><br>It is ignored if `–w ins` | Integer that specifies the contig number that will have the causal mutation. |
| `"PositionCausalMutation"`<br><br>It is ignored if `–w ins` | Integer with the position, in the contig specified with `"ContigCausalMutation"`, of the causal mutation defined in base pairs. It must be between 1 and the length of the contig. If `-w snp` and the position does not correspond to a G or C, Easymap v.2 will use the closest G or C instead to generate an EMS-like mutation. |
| `"NumberRecombinantChromosomes"`<br><br>It is ignored if `–w ins` | Integer that represents the number of haploid $F_2/M_2$ recombinant genomes to create. |
| `"RecombinationFrequencies"`<br><br>It is ignored if `–w ins` | Recombination frequency distribution of each contig provided in `-r`. It must contain a `“name”:“value”` pair for each contig in the reference sequence. `“value”` is a list of `event,frequency` pairs, where event is the number of crossover points per contig and frequency is the percentage of contigs in a recombinant contig population. `event,frequency` pairs are separated by a dash. `“name”:“value”` pairs are separated by commas. |
| `"Library"` | Type of fragment library for sequencing. Choose between `se` (single-end library) and `pe` (paired-end library). |
| `"FragmentSize"` | Average size of the library fragments. It must be an integer equal to or greater than the read length specified in `"ReadSize"`. |
| `"FragmentSd"` | Standard deviation of the size of the library fragments. It must be a positive integer. |
| `"ReadSize"` | Average size of the reads. It must be a positive integer. |
| `"ReadSd"` | Standard deviation of the size of the reads. It must be a positive integer. |
| `"ReadDepth"` | Number of times in average that each nucleotide in the reference sequence is read. It must be a positive integer. |
| `"ErrorRate"` | Integer in the interval [0–5] that represents the base-calling error rate in percent. |

| | |
|---|---|
| `"GCbias"` | Integer in the interval [0–100] that represents the GC content bias strength of the library. Setting this to >0 penalizes the creation of fragments with non-neutral GC content. The probability that the genomic sequence is present in the set of reads created is inversely proportional to the bias of the GC content and to the strength value set by the user. |

#### XIII. Appendix D: Other installation setups

This section includes some helpful notes for the installation of Easymap v.2 in different setups. Please note that the performance of virtual machines and apps is reduced compared to that of a direct installation of an OS, so for low-performance machines these options are unadvisable.

**Installation in Windows 10 Ubuntu app.** After installing Ubuntu from the Microsoft Store (search for “Ubuntu 18.04 LTS”), the user may be prompted to enable the “Windows Subsystem for Linux” feature of the Windows 10 OS. To do so, run the Windows 10 PowerShell as administrator, type the following command on the PowerShell, hit enter and then restart the system:

```
Enable-WindowsOptionalFeature -Online -FeatureName Microsoft-Windows-Subsystem-Linux
```

Once the Ubuntu app is working, Easymap v.2 can be installed by running the installation script as explained in section II or in the Quickstart Installation Guide. Make sure the dependencies specified in Appendix A are installed by running the commands provided. Automated startup of the Easymap v.2 dedicated server does not work within this environment; to access the web interface, the server needs to be started manually after each reboot with the following commands:

```
$ cd /path/to/easymap.v2
$ bash launch-server.sh <port-number>
```

The interface is accessed at http://localhost:<port-number> and can only be used locally; remote access to the server is restricted by the system.

**Installation in Amazon EC2.** For remote installation in a cloud computing service, the remote instance should be set up to allow HTTP access. In Amazon EC2, the recommended Amazon Machine Image (AMI) is Ubuntu Server 18.04. During setup, in the “Configure Security Group” tab, click on “Add Rule”, choose the “All traffic” option, and change the “Source” option to “Anywhere”. Once the instance is set up, Easymap v.2 is installed as explained in section II. To access the web interface, point the browser to the Public DNS address assigned to the instance, adding “:<port-number>” at the end.

**Installation in Oracle Virtualbox.** Oracle Virtualbox is a free software for running virtual machines available for Windows and Mac OS. Ubuntu Server can be downloaded for free from https://releases.ubuntu.com/18.04.4/ubuntu-18.04.4-live-server-amd64.iso and installed within the Oracle Virtualbox environment. This link contains general instructions for the installation of the Ubuntu OS in a Virtualbox virtual machine: https://linuxhint.com/install_ubuntu_virtualbox_2004/. During the configuration of the virtual machine, the network should be set as “bridged network” so that the web interface can be accessed later. A minimum of 4 Gb of RAM and a fixed-size virtual hard drive of at least twice the size of the reads to be processed is recommended for the virtual machine. Once the virtual machine is running, access the Ubuntu terminal and install Easymap v.2 as explained in section II or in the Quickstart Installation Guide. After installation, access the Easymap v.2 web interface using the virtual machines IP address from the web browser at: http://<ip-address>:<port-number>. To find the IP address, run the following command and select the address from the “inet” field:

```
$ ip addr show
```

Output example: inet 11.22.33.44/55 brd 10.1.31.255 scope global dynamic
IP address: 11.22.33.44

#### XIV. Appendix E: Modifying Easymap v.2 workflows

Easymap v.2 was designed to be as user-friendly as possible, hardcoding most of the parameters used during the alignment of the reads, the variant-calling or posterior filtering steps, to save the user from having to set any complex parameters. Advanced users can adapt Easymap v.2 workflows to any specific need or add steps to the analyses. The main workflows used by Easymap v.2 are located in the folder `/easymap.v2/workflows`. The code is written in commented and well-organized blocks to easily find and isolate any specific part of the analysis. The parameters used for the read alignment and variant calling steps can be modified in the Samtools, Hisat2, and BCFtools commands.

### Easymap v.2 Quickstart Installation Guide

This is a simplified guide for **Easymap v.2** installation in the main supported Operating Systems. Check the **Easymap v.2 Documentation** at http://genetics.edu.umh.es/resources/easymap/ for more detailed information and other installation setups. **System administrator privileges and internet access** are required for the installation. The following steps are meant to simplify the installation; advanced users can alter the process for specific needs.

#### 1 Installation Environment

Easymap v.2 is meant to be installed within a **Linux system**, these are a few setups that can be used:

- A physical machine running **Ubuntu**, **Linux AMI** or **Red Hat**
- The **Ubuntu 18.04 app**, available for download in the **Windows 10 Microsoft Store**. *You may be prompted to enable the "Windows Subsystem for Linux" feature of the Windows 10 OS. To do so, run the Windows PowerShell as administrator, run the following command and restart your system:*

```
Enable-WindowsOptionalFeature -Online -FeatureName Microsoft-Windows-Subsystem-Linux
```

- A virtual machine (VM) running Linux within **Windows** or **Mac OS**. *Get the Ubuntu ISO at https://releases.ubuntu.com/18.04.4/ubuntu-18.04.4-live-server-amd64.iso and mount it using Oracle Virtualbox. Performance in virtual machines is limited but can be sufficient. Choose a "Bridged connection" while setting up the virtual machine to be able to access the graphic interface later.*

#### 2 Easymap v.2 Installation

Open the Linux console and run the following commands (ignoring $ symbol):

```
$ cd ~                                                              Moves to home folder

$ wget http://genetics.umh.es/other_files/genetica%20umh%20es       Downloads the Easymap
       /Easymap/easymap-installer-v2.sh                             v.2 installer

$ sudo bash easymap-installer-v2.sh                                 Installs Easymap v.2
```

Select your operating system from the interactive menu and wait for the installer to finish. You will get a "Easymap successfully installed" message at the end. Installation can take up to 30 minutes. *In the unlikely case your machine doesn't have wget installed, use the command "*`sudo apt install wget`*" in Ubuntu or "*`sudo yum install wget`*" in other distributions.*

#### 3 Access Easymap v.2

- For **local** access, point your web browser to **http://localhost:8100**
- For **remote** access (from another machine or VM), point your web browser to **http://<IP-address>:8100**

#### Additional notes

- See Easymap v.2 Documentation for further details and other installation setups.
- Manual server rebooting may occasionally be necessary. If at any time you cant access the graphic interface please run the following commands in the console:

```
$ cd ~/easymap.v2                              Moves to easymap v.2 directory

$ bash launch-server.sh <port-number>          Launches easymap v.2 server
```

**Important**: You must replace `<port-number>` with any number "81XX" between 8101 and 8200 and use the same port number to access easymap ( http://localhost:81XX ).

- A preview version of the graphical interface is available for testing at: http://atlas.umh.es/easymapv2

**Supplementary Table S1.** Validation of Easymap v.2 for variant density mapping analyses

| Data source | Reference | Species | Mapping strategy | Mutant | Project name in the preview Easymap v.2 interface |
|---|---|---|---|---|---|
| Provided by the authors | Zuryn *et al*. (2010) | *C. elegans* | Backcross | *mutA* | 2020-08-27-13:19:26_mutA |
| | | | | *mutD* | 2020-08-27-17:07:57_mutD |
| | | | | *mutH* | 2020-08-27-20:56:44_mutH |
| Provided by the authors | Zuryn *et al*. (2014) | *C. elegans* | Backcross | *jmjd-3.1* | 2020-08-29-18:34:13_37 |
| | | | | *egl-27* | 2020-08-28-20:38:40_83 |
| Provided by the authors | Svensk *et al*. (2016) | *C. elegans* | Backcross | *sma-1* | 2020-09-08-14:04:40_40 |
| | | | | *dpy-23* | 2020-09-08-20:59:14_41 |
| | | | | *et43* | 2020-09-09-09:07:16_43 |
| PRJNA476333 | Klein *et al*. (2018) | *Z. mays* | Outcross | *ten* | 2020-08-27-10:08:18_TEN |

**Supplementary Table S2.** Validation of Easymap v.2 for QTL-seq mapping analyses

| Data source | Reference | Species, cultivar | Mapping population | Trait, mutation, or QTL | Project name in the preview Easymap v.2 interface |
|---|---|---|---|---|---|
| Requested from the authors | Illa-Berenguer *et al*. (2015) | *S. lycopersicum* | $F_2$ | *fw11.2* | 2021-11-06-11:59:05_139 |
| | | | | *lcn2.4*, *fw3.3*, *lcn5.1*, *lcn6.1* | 2021-11-13-17:39:12_75 |
| | | | | *fw1.1* | 2021-11-06-11:59:19_143 |
| DRA001007 | Fekih *et al*. (2013) | *O. sativa*, cv. Hitomebore | $M_3$ | Hit11440 | 2021-11-06-11:59:30_Hit11440 |
| | | | | Hit9188 | 2021-11-06-11:59:35_Hit9188 |
| PRJDB4643 | Hisano *et al*. (2017) | *H. vulgare* | Double haploid | *blp* | 2021-11-06-11:59:25_blp |
| | | | | Net blotch resistance | 2021-11-06-11:59:40_nbr |
| DRA000809 | Takagi *et al*. (2013) | *O. sativa*, cv. Dunghan Shali | $F_2$ | Seedling vigor | 2021-11-06-11:59:54_OsDung |
| | | *O. sativa*, cv. Nortai | RIL | Blast fungus resistance | 2021-11-06-12:00:04_OsNor |
| | | *O. sativa*, cv. Arroz da terra | RIL | Germination rate at low temperatures | 2021-11-06-11:59:48_OsArroz |
| | | *O. sativa*, cv. Iwate96 | RIL | Grain amylose content | 2021-11-06-11:59:56_OsIwa |
| SRR6831103, SRR6831101 | Wang *et al*. (2018) | *O. sativa*, cv. Nipponbare | $F_2$ | *wb1* | 2021-11-06-12:00:40_Wang18 |
| SRR5739122, SRR5739123 | Yang *et al*. (2017) | *O. sativa*, cv. Nipponbare | $F_2$ | *qNUE6* | 2021-11-30-18:17:01_Yang2017 |
